# Molecular characteristics of chronic hepatitis B virus infection among voluntary blood donors in Kinshasa, Democratic Republic of the Congo

**DOI:** 10.1101/740803

**Authors:** Pati Moloko Maindo, Didier Anzenza Mudwahefa, Jean-Pierre Kambala Mukendi, Sylvain Ramazani Yuma, Jérémie Masidi Muwonga, Gerald Misinzo

## Abstract

**Background:** Hepatitis B represents a major global health problem. Despite the high endemicity of hepatitis B in Sub-Saharan Africa, little is known about the molecular characteristics of chronic hepatitis B virus (HBV) infection in Africa, and there are very few published studies that describe the genetic characteristics of HBV in asymptomatic adults in DRC. The present study aimed at determining the molecular diversity of chronic HBV infection in voluntary blood donors in Kinshasa, DRC.

**Methods:** Blood samples from 582 voluntary blood donors at the National Blood Transfusion Centre in Kinshasa, DRC, were screened for hepatitis B surface antigen (HBsAg) using enzyme-linked immunosorbent assay (ELISA). Partial amplification and sequencing of S gene in HBV-positive samples was conducted.

**Results:** The presence of HBsAg was detected in 6.9 % (40/582) blood donors. Phylogenetic analysis based on partial S gene nucleotide sequences of HBV showed that the majority (66.7 %, 10/15) of HBV strains clustered into genotype A, followed by the genotypes E (26.6 %, 4/15) and D (6.7 %, 1/15). Genotype A strains were classified into subgenotype A1, quasi-subgenotype A3, and subgenotype A4, with quasi-subgenotype A3 being predominant. One new genotype A strain did not cluster with any existing HBV/A subgenotype or quasi-subgenotype.

**Conclusions:** The present study highlights the high genetic variability of chronic HBV infection in DRC, and the possibility of a new HBV/A subgenotype, suggesting that HBV has a long evolutionary history in DRC. Further molecular characterization of complete HBV genomes is needed for a more accurate assessment of HBV genetic variability and its clinical significance in DRC, as partial sequences are not appropriate for determining HBV new subgenotypes.

## Introduction

Hepatitis B virus (HBV) infection represents a major global health problem. According to the World Health Organization, an estimated 257 million people live with chronic hepatitis B infection [1], which is characterized by the persistence of hepatitis B surface antigen (HBsAg) in blood for a period of more than 6 months [2]. About 15 to 40 % of HBV-infected patients develop cirrhosis, liver failure, or hepatocellular carcinoma (HCC) [3, 4]. In 2015, about 887 000 people died from end-stage liver disease or HCC [1]. The global prevalence of chronic HBV infection, defined as the prevalence of HBsAg, varies widely worldwide. Sub-saharan Africa have the highest rates, since more than 8 % of the population in these regions have chronic HBV infection [5].

Owing to the high genetic variability of HBV, eight confirmed HBV genotypes designated A to H, two tentative genotypes named I and J, and around forty sub-genotypes have been described [6]. The HBV genotypes and subgenotypes are characterized by a minimum sequence divergence of 7.5 and 4 % of the entire genome, respectively [6–8]. Recently, some subgenotypes have been redefined into quasi-subgenotypes. Quasi-subgenotypes are lineages, which during full analysis taking into account all circulating strains of the same genotype and subgenotypes, do not fit the criteria of the subgenotypes [9]. Different HBV genotypes and subgenotypes/quasi-subgenotypes display differences in transmission routes, including sexual intercourse, unsafe injections, blood transfusions and mother-to-neonate transmission [6]. Moreover, many studies have shown that some genotypes are associated with particular prognoses, such as progression to HCC, acute forms of hepatitis B or response to antiviral treatment. Thus, the knowledge of HBV genotypes and subgenotypes can help in predicting clinical outcome and planning suitable treatment [6, 10–13]. In addition, according to geographical distribution, HBV genotypes vary between continents, geographical regions, countries, and even between geographical regions within countries [12]. Genotypes A and E prevail in Africa, the former being mainly found in Eastern, Central and Southern Africa, and the latter being particularly restricted to West and Central Africa [13, 8]. Genotype D is predominant in Northern Africa [13].

Yet, data on the molecular characteristics of HBV in adults living in DRC are very limited. Some studies have been performed on very small sample sizes of adults [14–16]. Other studies included either symptomatic patients with jaundice from different provinces of the country [17] or adults living in the eastern part of DRC [18, 19]. The present study was performed in asymptomatic adults in Kinshasa, the capital city of DRC, located in the western part of the country.

## Materials and methods

### Study area, study design and sample size

The present cross-sectional study was conducted in Kinshasa, the capital city of DRC that lies on longitude 15° 19’ 16’’ East of Greenwich and latitude 4° 19’ 19’’ South of Equator [20]. A total of 582 voluntary blood donors were enrolled in the present study between November 2014 and December 2014.

### Study population and study exclusion criteria

Blood samples from blood donors collected by the National Blood Transfusion Centre (CNTS) during the study period were included in the present study. According to the CNTS blood donation criteria, blood donors who had a history of transfusion or jaundice, and all those belonging to a high-risk group, including drug addicts, professional sex workers and those with multiple sexual partners, were excluded from blood donation.

### Detection of HBsAg

Serum samples were routinely screened for HBsAg using ELISA (Hepanostika^®^ HBs, Biomérieux, France and Abbott GmbH & Co. KG, Wiesbaden, Germany), following manufacturer’s instructions. Then, blood from HBsAg positive samples (50 µL) was drawn from either ethylene diamine tetra acetic acid (EDTA) vacutainer tubes within 72 hours, or corresponding blood bags containing citrate phosphate dextrose adenine (CPDA) anticoagulant within 7 days following blood donation, and spotted into cards (FTA^®^ classic cards Whatman^®^ 3 mm). Afterwards, FTA classic cards were dried and stored at room temperature until used for deoxyribonucleic acid (DNA) detection and sequencing.

### Detection of HBV DNA

FTA paper circles of 0.5 mm diameter containing blood samples were suspended in 200 µL of lysis buffer and digested with 20 µL of proteinase K for 5 hours at 55 °C, followed by a standard nucleic acid extraction method using QIAamp genomic DNA Mini Kit (Qiagen Sciences, Maryland, USA), following the manufacturer’s instructions. Then, HBV DNA detection was performed by amplification of the partial S-gene, using a nested polymerase chain reaction (PCR) that was expected to yield a product of approximately 400 base pairs. The following primer combinations were used: HBV_S1F (5’ - CTA GGA CCC CTG CTC GTG TT - 3’, nucleotide position 179) with HBV_S1R (5’ - CG AAC CAC TGA ACA AAT GGC ACT - 3’, nucleotide position 704) as outer primers, and HBV_SNF (5’ - GTT GAC AAG AAT CCT CAC AAT ACC - 3’, nucleotide position 217) and HBV_SNR (5’-GA GGC CCA CTC CCA TA −3’, nucleotide position 658) as inner primers. These primers were previously used by Forbi *et al.* [21], with some modifications. After an initial denaturation step (10 min at 95 °C), DNA amplification was performed for 40 cycles at 95 °C for 30 sec, 55 °C for 45 sec and 72 °C for 45 sec, and a final extension at 72 °C for 10 minutes using a Veriti 96-Well Thermal Cycler (Applied Biosystems, Jurong, Singapore). The first PCR round was performed with 2 μL of DNA and 1 U of *Taq* DNA polymerase (Invitrogen, Carlsbad, CA, USA) in a final volume of 20 μL. The second PCR round was performed with 2 μL of DNA of the PCR product from the first round of amplification and 1 U of *Taq* DNA polymerase (Invitrogen, Carlsbad, CA, USA) in a final volume of 20 μL.

### Sequencing of PCR products

A total of 5 μL of PCR amplicons were loaded onto a 1.5 % GelRed-stained agarose and the DNA bands were visualized using a gel documentation system (Gel Doc Ez Imager, Bio-Rad, CA, USA). PCR products were denatured at 95 °C for 10 min in the thermal cycler to avoid degradation by nucleases. Then, they were sealed and stored at −40 °C until sequencing. The sequencing was carried out by Macrogen Incorporation (Macrogen Europe, Amsterdam, The Netherlands) by using the automated 3730xl DNA Analyzer machine and BigDye XTerminator Cycle Sequencing Kit version 3.0 (Applied Biosystems, Foster City, CA, USA). HBV_SNF and HBV_SNR were used as primers for sequencing of the partial S gene.

### Phylogenetic analyses

The nucleotide sequences obtained were submitted to GenBank and were given the accession numbers from KR535608 to KR535622 (Table S1). A similarity search at GenBank nucleotide database was conducted using the Basic Local Alignment Search Tool for nucleotides (BLASTn) [22]. The new HBV sequences from Kinshasa along with 64 representative strains of genotypes A to H (Fig.1) were aligned using Clustal W, and a phylogenetic tree was constructed using the Molecular Evolutionary Genetics Analysis software (MEGA6.05) by the Maximum Likelihood method and the Tamura three-parameter model of nucleotide substitution [23]. Bootstrap testing of phylogeny was inferred following 1000 replications to assess the reliability of the clusters. Values equal to or greater than 50 were indicated on the branches.

**Figure 1.**
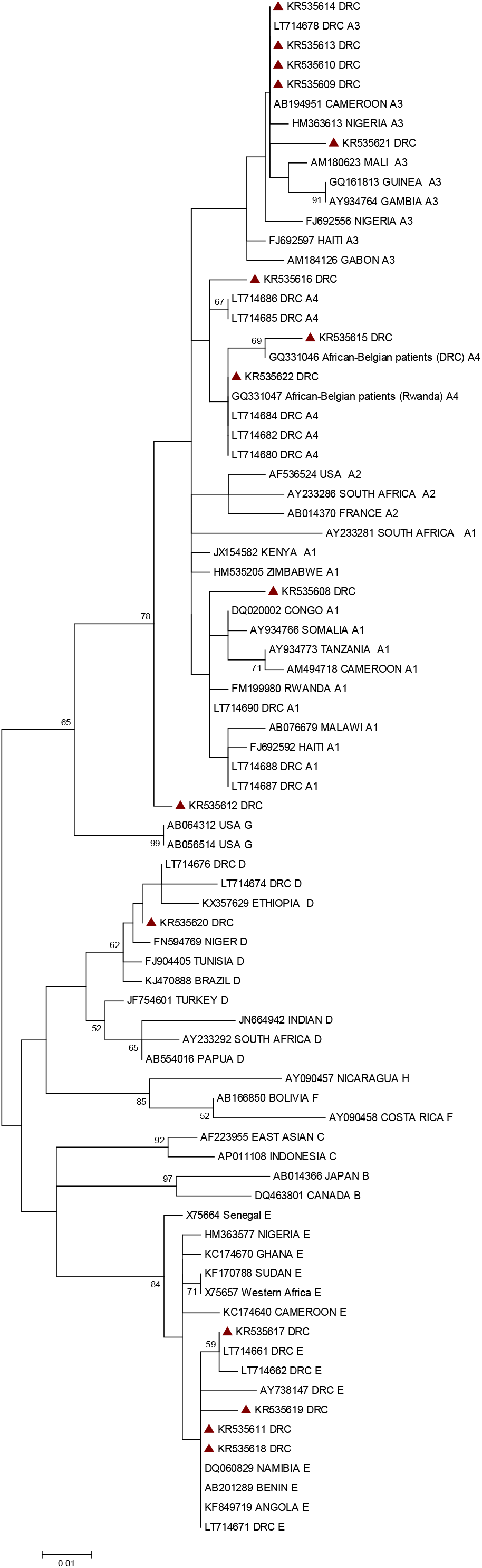
Genetic relationship of HBV in blood donors of Kinshasa. Maximum Likelihood method was used to infer phylogenetic relationship, with the Tamura 3-parameter nucleotide substitution model and bootstrapping of 1000 replicates as implemented in MEGA6.05 software. Relevant bootstrap values, using a cut-off value of 50 %, are shown. Sequences retrieved from GenBank are represented by their accession and country names. New HBV-DRC strains are preceeded by a filled red triangle. The scale bar indicates the number of nucleotide substitutions per site.(PDF 14 kb)

## Results

In the present study, the presence of HBsAg was detected in blood donors, using ELISA. A total of 582 blood donors including 503 men (86.4 %) and 79 (13.6 %) women were enrolled. Among the 582 blood donors, 40 (6.9 %) were positive for HBsAg. The median age of blood donors was 28 years (ranged from 18-64 years). Out of the 40 HBsAg-positive samples, 36 were subjected to HBV-DNA extraction. The four other samples could not be traced in the freezers after an initial HBsAg screening. Hepatitis B virus DNA was also extracted from four samples used as HBsAg positive controls by the CNTS. These positive controls were samples from blood donors, which were strongly positive in HBsAg serological detection and were then used as positive controls to subsequent ELISA. The detection of HBV DNA was successful in 50% (18/36) of HBsAg positive samples and 2/4 of HBsAg positive controls. From the 18 DNA samples, 14 were selected for sequencing, based on quality of PCR products. Out of the 14 PCR products subjected to nucleotide sequencing, 13 were successfully sequenced. Both DNA detected in the two positive controls were also successfully sequenced. Thus, a total of 15 nucleotide sequences were obtained and were named beginning with “DRC-Kin” followed by study identification numbers or letters, e.g. DRC-Kin-9126519 or using their accession numbers (Table S1). Ten (66.7 %) out of the 15 new nucleotide sequences clustered into genotype A. Four new nucleotide sequences (26.6 %) clustered into genotype E. One (6.7%) nucleotide sequence clustered into genotype D. The new HBV/A sequences belonged to quasi-subgenotype A3 (5/10), subgenotype A4 (3/10) and subgenotype A1 (1/10). Majority of the new HBV strains were closely related either to strains from DRC alone, or to strains from DRC and other African countries. One of the new HBV/A sequences (KR535612 DRC) did not cluster with any HBV/A subgenotype (Fig. 1).

## Discussion

The present study aimed at determining HBV genotypes among voluntary blood donors in Kinshasa, the capital city of DRC, whose inhabitants originate from different provinces of DRC. The high diversity of genotypes we found (genotypes A, E, and D) is consistent with results reported by previous studies performed in samples from subjects originating from different provinces of DRC [17] and in 6 samples from Kinshasa and the western part of DRC [14].

Majority of HBV strains found in the present study (66.7%) belonged to genotype A, followed by genotype E (26.6%). Though Kinshasa is located in the western part of DRC, our results seem to be contradictory to earlier studies reporting the predominance of HBV/E in the western part of DRC [14] and in samples collected from subjects having acute febrile jaundice and originating from different provinces of DRC [17]. This discrepancy may be due to differences in study populations and/or population sizes. Kinshasa is also likely to be an exception in the West-African genotype E crescent, but this has to be confirmed by future studies performed in larger sample sizes.

Seven HBV/A subgenotypes were described, namely HBV/A1, HBV/A2, HBV/A3, HBV/A4, HBV/A5, HBV/A6 and HBV/A7 [2428, 2529]. But further, a new classification was proposed, grouping together subgenotypes A3, A4, A5 and A7 into “quasi-subgenotype A3” and renaming HBV/A6 into HBV/A4 [9, 2529]. The new HBV/A strains we reported clustered into three subgenotypes, namely A1, A4 and quasi-subgenotype A3, the latter being the most frequent HBV/A subgenotype. This supports previous studies reporting HBV/A3 as predominant in West/Central Africa [26] The particularity of HBV/A4 is to be a recent HBV/A subgenotype that has so far been reported in very few Africa countries including DRC and two out of its neighbouring countries, namely Republic of Congo and Rwanda [16, 27]. This high HBV/A variability suggests that HBV has a long evolutionary history in Kinshasa, DRC. One HBV/A new strain study did not cluster with any known HBV/A subgenotype. The possibility that this strain be a new HBV/A subgenotype is not excluded. Further molecular characterization of complete HBV nucleotide sequences is needed in order to accurately assess the HBV/A genetic variability in DRC, as results obtained from the partial sequence are not appropriate for introducing novel HBV subgenotypes [9, 28].

## Conclusion

The present study principally aimed at determining the HBV genotypes circulating in asymptomatic adults in Kinshasa, DRC. Three HBV genotypes were found, namely A, D and E, with a predominance of genotype A/ quasi-subgenotype A3. One HBV/A strain clustered into none of the known HBV/A subgenotype. Further molecular characterization of complete HBV nucleotide sequences is needed for a more accurate assessing of HBV genetic variability and its clinical significance in DRC.

## Additional files

**Additional file 1: Table S1.** HBV strains sequenced in the present study. (DOC 13 kb)

## Abbreviations

BLASTn: Basic Local Alignment Search Tool for nucleotides
CNTS: National Blood Transfusion Centre
CPDA: citrate phosphate dextrose adenine
DNA: deoxyribonucleic acid
DRC: Democratic Republic of the Congo
EDTA: ethylene diamine tetra acetic acid
ELISA: enzyme-linked immunosorbent assay
HBsAg: hepatitis B surface antigen
HBV: hepatitis B virus
HCC: hepatocellular carcinoma
MEGA: Molecular Evolutionary Genetics Analysis software
PCR: polymerase chain reaction

## Acknowledgments

We appreciate the excellent technical assistance of Christopher Kasanga from Sokoine University of Agriculture, Morogoro, Tanzania, Donatien Kayembe and Mireille Nganga from the University Clinics of Kinshasa, Kinshasa, DRC, of Paul Masiangi from the School of Public Health in Kinshasa, DRC, and of Eddy Sokolua from the CNTS, Kinshasa, DRC.

## Availability of data and materials

The datasets generated and/or analysed during the current study are available in GenBank with accession numbers KR535611-KR535622.

## Authors’ contributions

PMM and GM and conceived and designed the experiments. PMM, GM and JPKM performed the experiments. GM, PMM and DAM analyzed the data. GM, PMM, SRY and DAM contributed reagents/materials/analysis tools. GM, PMM and JPKM drafted the paper. All authors read and approved the final manuscript.

## Consent for publication

Not applicable.

## Ethics approval and consent to participate

All blood donors were older than 18 years and provided informed consent to participate. The approval and ethical clearance to carry out this study was granted by the ethics review committee of the Public Health School of DRC (ESP/CE/027/2015).

